# Deregulation of Y-linked protamine-like genes in sex chromosome-biased spermatid demise

**DOI:** 10.1101/2022.12.04.519051

**Authors:** Jun I. Park, George W. Bell, Yukiko M. Yamashita

## Abstract

Meiotic drive is a phenomenon wherein a genetic element achieves a higher rate of transmission than dictated by Mendelian segregation (*1-3*). One proposed mechanism for meiotic drivers to achieve biased transmission is by sabotaging essential processes of gametogenesis (e.g. spermatogenesis), leading to demise of gametes that contain their opponents (*1*). Studies in *D. simulans* have recently found that a set of meiotic driver genes contain a sequence homologous to protamines (*4, 5*), critical proteins that package sperm chromatin (*6-8*). However, the underlying mechanisms of drive and the relevance of protamine-like sequences in meiotic drive remain unknown. While studying the function of Modulo, the homolog of Nucleolin in *Drosophila melanogaster* (*9, 10*), we unexpectedly discovered Y-linked protamine genes function as a meiotic driver: we found that *modulo* mutant’s known sterility is caused by deregulation of the autosomal protamine-like gene (*Mst77F*) and its Y chromosome-linked homolog (*Mst77Y*). Modulo regulates these genes at the step of polyadenylation of the transcripts. We show that *Mst77Y* likely acts as a dominant-negative form of *Mst77F*, interfering with the process of histone-to-protamine transition, leading to nuclear decompaction. Overexpression of *Mst77Y* in a wild-type background is sufficient to cause nuclear decompaction and results in the biased demise of X chromosome-bearing sperm. We propose that dominant-negative protamine variants may be a common strategy found in male meiotic drive and may explain known rapid divergence of protamine genes.

**Significance statement:** Protamines are small, highly positively charged proteins that are required for packaging DNA to produce mature sperm with highly-condensed nuclei capable of fertilization. Even small changes in the dosage of protamines in humans is associated with infertility. Yet, despite their essential function, protamines are rapidly evolving. It has been speculated that protamines’ rapid divergence may be explained by their potential participation in genomic conflict. Our work implicates the involvement of Y chromosome-linked multicopy protamine-like genes in meiotic drive in *Drosophila melanogaster*. Our results suggest that dominant negative protamines can sabotage the process of nuclear compaction during spermiogenesis, revealing a potential cellular mechanism of sperm killing in meiotic drive.

---

In many species, spermatids undergo the process of nuclear compaction, an essential process to produce sperm that are capable of fertilization (*8, 11*). Nuclear compaction is critical for the sperm’s hydrodynamic performance and protecting the paternal genome against mutagens (*12, 13*). Nuclear compaction involves the dramatic chromatin reorganization mediated by the histone-to-protamine transition (6, *8, 11-14*). Protamines are small, positively charged proteins that replace histone-based nucleosomes to achieve the extreme degree of DNA compaction often seen in sperm (*11*). These protamines are required for fertility across many different species. In humans, there are two main protamine genes (PRM1 and PRM2), and even small changes in the ratio between the amount of these protamines is correlated with subfertility(*11, 14*).

Although protamines are essential for fertility, they are rapidly evolving across many different species (*2, 12, 15*). In mammals, PRM1 and PRM2 are required for fertility in humans and mice, while PRM2 has become non-functional in bulls and boars (*12*). Likewise, closely related *Drosophila* species have independently evolved many different protamine-like genes (*2*). One possible explanation for the rapid evolution of these essential genes is their involvement in genetic conflict, such as meiotic drive, wherein drivers and suppressors coevolve in a genetic ‘arms race’ to overcome one another (*2, 4, 5*).

Interestingly, recent studies have shown a possible role for protamines in the Winters *sex-ratio* (*SR*) system of meiotic drive in *D. simulans* (*4, 5*). Normally, *distorter on the X (Dox*) and its paralogs on the X chromosome are repressed by autosomal small RNAs (*4, 5*). When *Dox* is derepressed, post-meiotic spermatids undergo “sperm killing”, where a subset of spermatids in a cyst become non-viable, and results in female-biased progeny, suggesting that X-containing sperm are favored (*16*). The cytological mechanism by which this “sperm killing” may occur has largely remained a mystery, although sperm killing is quite common in many systems of male meiotic drive (*1, 3*). Recently, it has been shown that *Dox* and its paralogs contain a large portion of the protamine gene (*4, 5*). However, whether there is a functional role for these protamine regions in *Dox* function is unknown.

While studying *D. melanogaster modulo* mutant, we discovered that *modulo* mutant misregulates the expression of protamine genes, leading to decreased incorporation of protamine Mst77F and increased incorporation of Mst77Y, the Y-linked homologs of Mst77F. Our data indicate that Mst77Y likely acts as a dominant negative form of Mst77F, sabotaging the process of histone-to-protamine transition. Interestingly, Mst77Y preferentially harms X-bearing spermatids, favoring the transmission of the Y chromosome, thereby acting as a meiotic driver. Our study provides mechanistic insights into how protamines participate in the process of meiotic drive and why protamine genes may be rapidly evolving.

## *modulo* mutant is defective in sperm nuclear compaction

Modulo is the *Drosophila* homolog of Nucleolin, a nucleolar protein implicated in RNA processing (*9*). Although *modulo* mutant males have been known to be sterile (*10, 17*), the cytological defects that lead to their sterility have not been characterized. We find that the *modulo* mutant exhibits defects in nuclear morphology transformation during late spermiogenesis. In wild type males, post-meiotic spermatid nuclei undergo well-documented morphological changes (*8*), from round spermatid stage, to ‘leaf’ stage, ‘canoe’ stage, resulting in highly condensed ‘needle’ stage nuclei, which is accompanied by the histone-to-protamine transition (Fig. 1A). Although *modulo* mutant germ cells proceeded through spermatogenesis normally until ‘canoe’ spermatid stage (Fig. 1B, C), the *modulo* mutant exhibited striking ‘decompaction’ of the nuclei after reaching the canoe stage (Fig. 1D, E). Immunofluorescence (IF) staining using anti-dsDNA, which has been previously used to assess the compaction status of nuclei (*18*), revealed that defective spermatid nuclei of *modulo* mutant are indeed decompacted (Fig 1F, G). Decompacting nuclei are initially TUNEL-negative (Fig. 1H, I), then become TUNEL-positive (Fig. 1J), suggesting that decompaction is not the result of cell death, but may rather be a cause. Overall, 100% of the *modulo* mutant testes exhibited nuclear decompaction phenotype (Fig. 1K), and it appeared that all nuclei eventually become decompacted and die, filling the distal end of the testis with cellular debris (Supplementary Fig. 1A, B). The eventual death of all sperm nuclei likely results in the entire lack of mature sperm in the seminal vesicles (Supplementary Fig. 1C, D) and the *modulo* mutant’s known sterility (*10, 17*).

**Figure 1.**
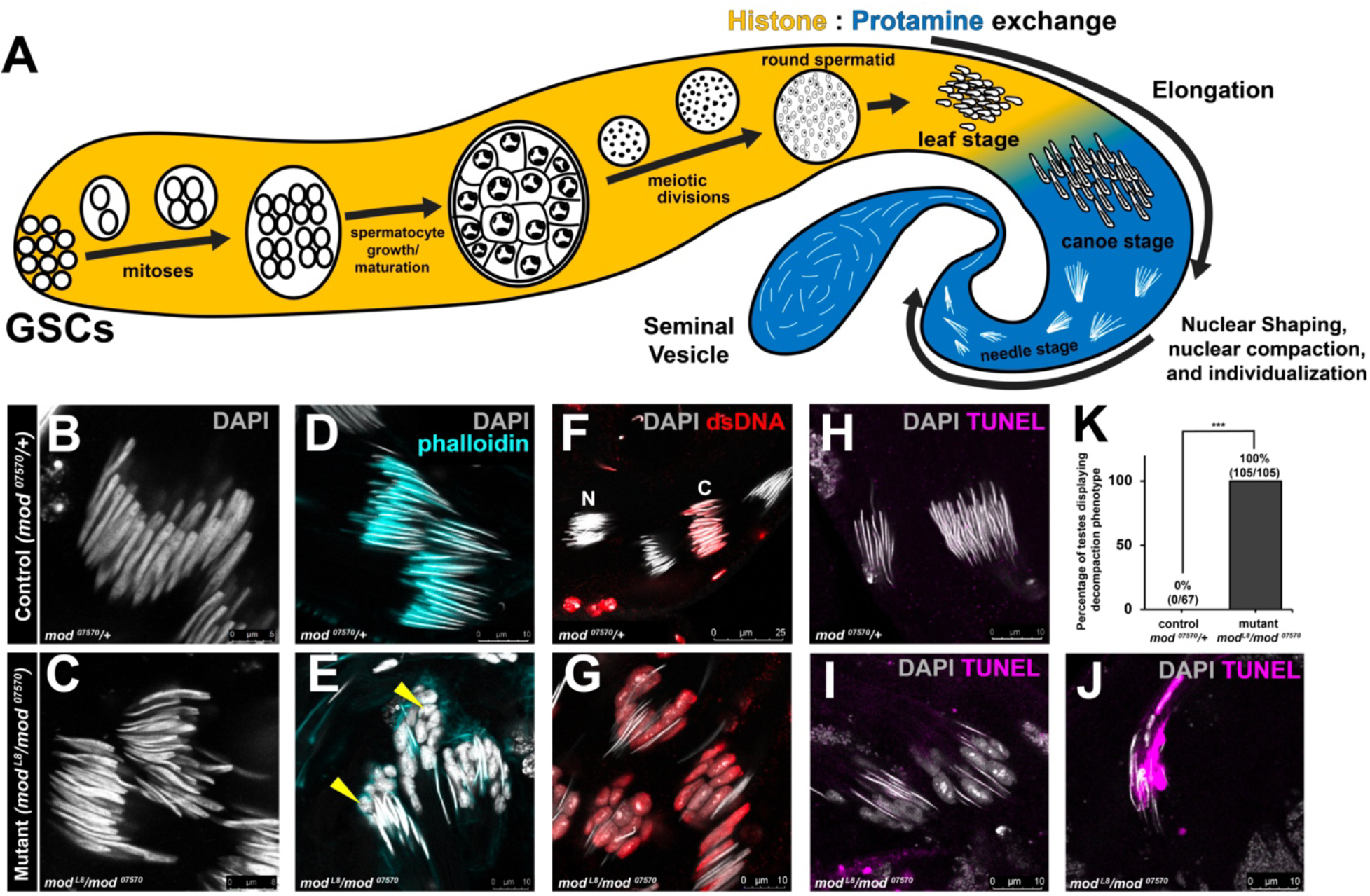
Sterility of *modulo* mutant is accompanied by defective spermatid chromatin compaction. A) Schematic of spermatogenesis in *Drosophila* proceeding from germline stem cells (GSCs) to mature sperm. Proceeding from meiotic divisions onwards, only nuclei are depicted. B, C) Representative images of canoe stage nuclei stained with DAPI (grey) in control male (*mod*^*07570*^*/+*) (B), and *modulo* mutant males (*mod*^*L8*^ / *mod*^*07570*^) (C). D, E) Representative images at the stage shortly before individualization stained with DAPI (grey) and phalloidin (cyan, to visualize the individualization complex) in control male (*mod*^*07570*^*/+*) (D), and *modulo* mutant males (*mod*^*L8*^ / *mod*^*07570*^) (E). Phalloidin staining shows relatively intact formation of individualization complex in *modulo* mutant, yellow arrowheads indicating decompacted nuclei. F, G) Representative images of late canoe to needle stages stained with anti-dsDNA (red) and DAPI (grey) in control (*mod*^*07570*^*/+*) (F), and *modulo* mutant males (*mod*^*L8*^ / *mod*^*07570*^) (G). N: needle-stage spermatids that do not stain for dsDNA due to advanced DNA compaction, C: canoe-stage spermatids that are less compact and positive for anti-dsDNA staining. H-J) TUNEL staining (magenta) of needle-shaped spermatid cysts in wildtype (*mod*^*07570*^*/+*) (H) and mutant (*mod*^*07570*^*/+*) males without (I) or with (J) TUNEL signal. K) Percentage of decompaction phenotype in *modulo* mutant vs wildtype males. *** indicates p<0.001 (unpaired Student’s t-test assuming unequal variances in 5 independent experiments). n (total number of testes counted per genotype) is presented on the bar graph.

## *modulo* mutant fails in histone-to-protamine transition

Because nuclear decompaction in *modulo* mutant occurs at stages when sperm chromatin is being reorganized by the histone-to-protamine transition, we explored if the *modulo* mutant is defective in this process. Histone-to-protamine transition occurs stepwise: 1) histone modification and removal, 2) transition protein incorporation then removal, 3) protamine incorporation (*8*). IF staining revealed that *modulo* mutant spermatids undergo proper histone removal and transition protein incorporation (Supplementary Fig. 2A-F), but fail to accumulate Protamines A/B and protamine-like protein Mst77F (Fig. 2A, B). Moreover, using a specific antibody (Supplementary Fig. 3A, B), we found that Mst77Y, Y-linked multicopy homologs of Mst77F (*19, 20*) (Supplementary Fig. 4A), strongly accumulated in *modulo* mutant spermatid nuclei, whereas it was barely detectable in control (Fig. 2C-G), suggesting that Mst77Y is aberrantly expressed in the *modulo* mutant. As the deletion of Mst77F and Protamine A/B does not cause nuclear decompaction as severe as *modulo* mutant (*21*), we infer that the incorporation of Mst77Y (in addition to depletion of Mst77F and Protamine A/B) causes the observed catastrophic nuclear decompaction seen in the all *modulo* mutant spermatids.

**Figure 2.**
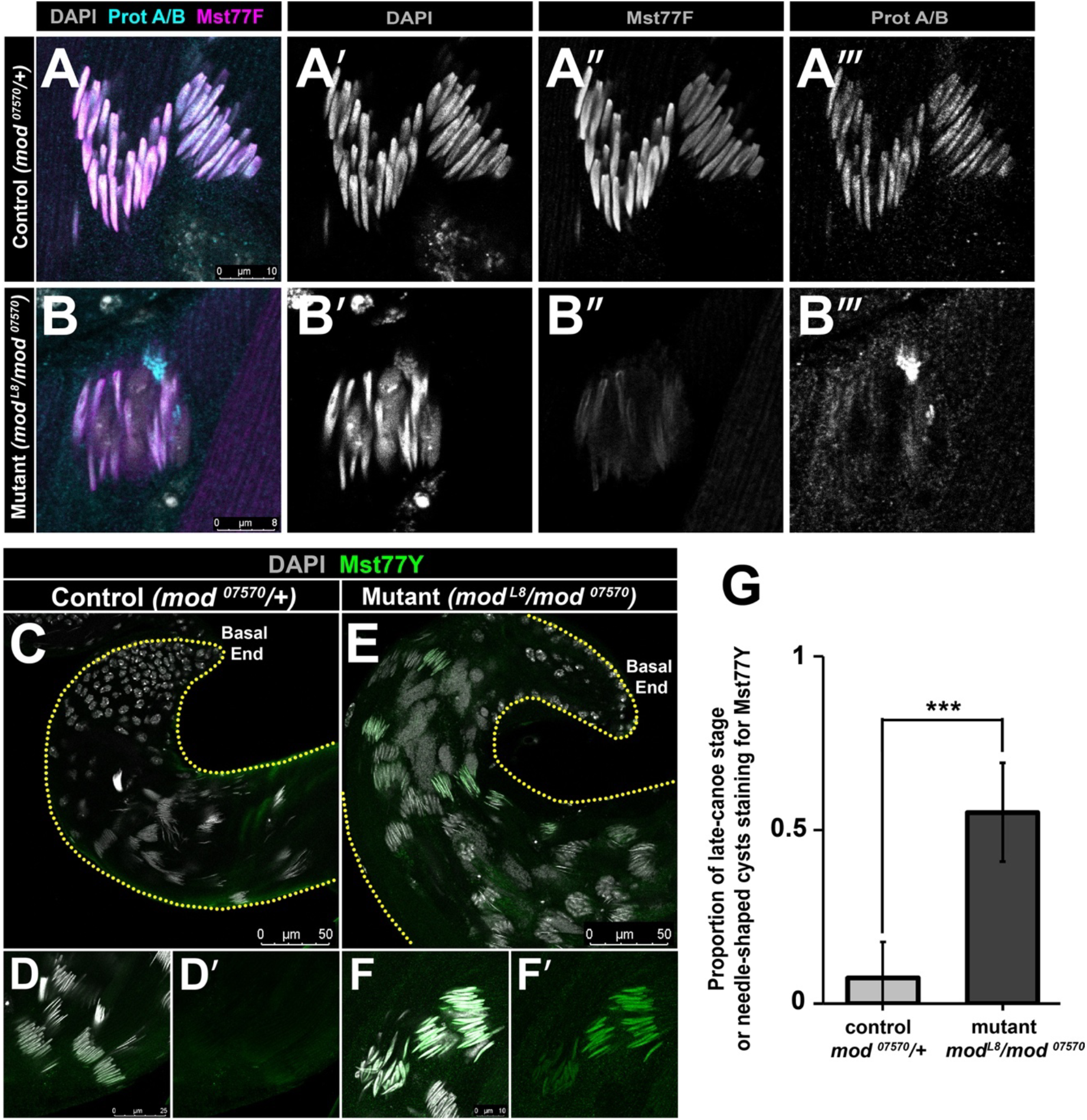
Nuclear decompaction in *modulo* mutant spermatids is associated with decreased incorporation of Mst77F and increased incorporation Mst77Y. A, B) Representative images of late canoe stage nuclei stained with DAPI (grey), anti-Prot A/B (cyan) and anti-Mst77F (magenta) in control (*mod*^*07570*^/+) (A) and mutant (*mod*^*L8*^/*mod*^*07570*^) (B) males. C-F) Representative images of canoe stage and needle stage spermatids at basal end of testis stained with DAPI (grey) and anti-Mst77Y (green) in control (*mod*^*07570*^/+) (C, D) and mutant *mod*^*L8*^/*mod*^*07570*^) (E, F) males. Dotted lines outline the testis. G) Proportion of canoe stage cysts with nuclei positive for Mst77Y staining in mutant (*mod*^*L8*^/*mod*^*07570*^) vs control (*mod*^*07570*^/+) males. *** indicates p ≤ 0.001 (unpaired Student’s t-test assuming unequal variances) with n=10 testes in control and n= 9 testes in mutant from 2 independent experiments. Exact p-values listed in Supplementary Table 1.

## Ectopic expression of *Mst77Y* alone is sufficient to cause nuclear decompaction

The *Mst77Y* genes have several interesting features. First, the gene locus contains 18 copies of *Mst77F* homolog (Supplementary Fig. 4A, B), which originated from a single event of *Mst77F* translocation to the Y chromosome, followed by gene amplification (*19, 20*). Second, many of the *Mst77Y* genes have mutations, which have resulted in changes in the position and number of critical arginine and lysine residues believed to be important for protamine function (*12, 22*). Other mutations have resulted in premature truncations (*19*). Note that anti-Mst77Y antibody was generated by using multiple peptides from Mst77Y that are distinct from Mst77F to increase specificity. The peptides were also designed to be able to identify all copies of Mst77Y, which feature similar mutations. Because *Mst77Y*’s mutations likely alter *Mst77F*’s normal function, we hypothesized that *Mst77Y* genes may function as a dominant-negative form of Mst77F. Accordingly, Mst77Y’s aberrantly high expression in *modulo* mutant may interfere with the process of normal histone-to-protamine transition.

To test the possibility that the expression of *Mst77Y* causes the nuclear decompaction phenotype, we generated transgenic lines that express *Mst77Y* under a male germline-specific *tubulin* promoter (*β2-tubulin* promoter) (*23-25*). From the 18 copies of *Mst77Y* homologs present on the Y chromosome (*19, 20*), we generated two lines expressing either *Mst77Y-12* (a full-length version) or *Mst77Y-3* (a truncated version due to premature stop codon) (Supplementary Figs. 4B, 5A-C), as the transcripts of these two genes have been previously detected by RT-qPCR (*19*). Strikingly, expression of either *Mst77Y-3* or *Mst77Y-12* (Supplementary Fig. 6A, B) recapitulated nuclear decompaction phenotype similar to that seen in *modulo* mutant (Fig. 3A-D): 45.7% and 43.2% of testes examined exhibited nuclear decompaction upon expression of *Mst77Y-3* or *Mst77Y-12*, respectively (Fig. 3E), suggesting that high *Mst77Y* expression is sufficient to cause nuclear decompaction in a subset of spermatids. Notably, in contrast to the eventual decompaction of all spermatids seen in the *modulo* mutant, *Mst77Y* overexpression alone does not cause sterility. We speculate that this might be due to the added effect of the decreased incorporation of Mst77F and Protamine A/B, in addition to high Mst77Y incorporation, seen in the *modulo* mutant.

**Figure 3.**
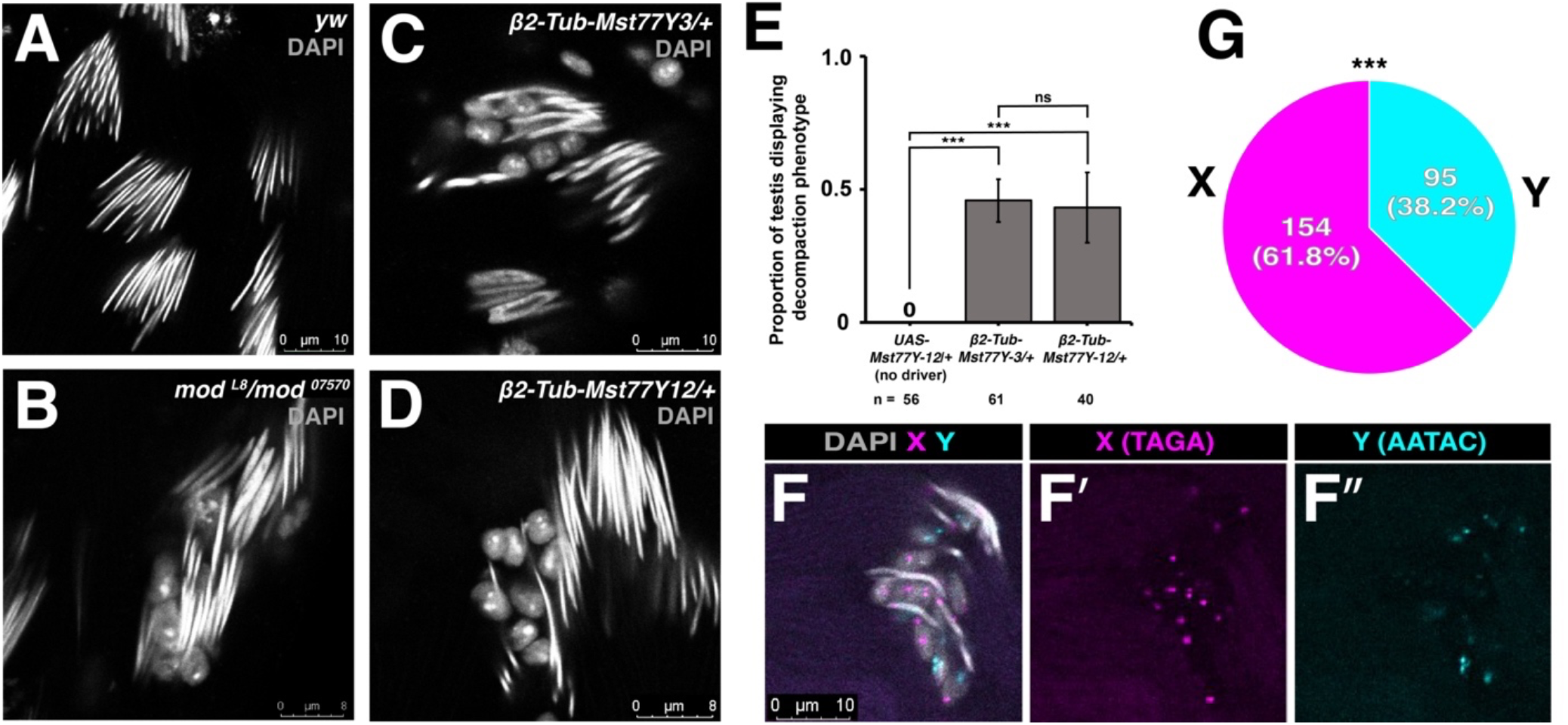
Mst77Y overexpression is sufficient to cause nuclear decompaction and causes biased decompaction of X chromosome-bearing spermatids. A-D) Representative images of needle-stage nuclei stained with DAPI (grey) showing normal morphology in control (*yw*) (A), and decompaction phenotype in *modulo* mutant (*mod*^*L8*^*/mod*^*07570*^) (B), transgenic males expressing Mst77Y-3 (truncated copy) (C) or Mst77Y-12 (full-length copy) (D) driven by *β2-tubulin* promoter. E) Proportion of testes displaying decompaction phenotype in transgenic Mst77Y males. Control (*UAS-Mst77Y12*/+) does not express Mst77Y12 due to the absence of driver. *** indicates p ≤ 0.001 (unpaired Student’s t-test assuming unequal variance) with n = 56 testes in control, n = 61 testes in *β2-tub-Mst77Y3/+* condition, n = 40 in *β2-tub-Mst77Y12/+* condition from 3 independent experiments. F) Representative images of DNA FISH of decompacted spermatids in Mst77Y-3 expressing males using AATAC-Cy5 (cyan, Y-specific probe) and TAGA-Cy3 (magenta, X-specific probe). G) Percentage of decompacted haploid nuclei containing X-chromosome vs Y-chromosome in Mst77Y3 expressing males. *** indicates P(X≥154) < 0.001 (exact Binomial Distribution) assuming p = 0.5 with n = 249 nuclei counted from 3 independent experiments. Exact p-values listed in Supplementary Table 1.

Considering that *Mst77F* overexpression does not cause such nuclear compaction defects as we observed with *Mst77Y* overexpression (*23*), we speculate that *Mst77Y* may act as a dominant-negative form, perhaps interfering with the function of *Mst77F*. This notion is further supported by the fact that a truncated version (*Mst77Y-3*) also causes the decompaction phenotype. Indeed, spermatid cysts of transgenic males expressing *Mst77Y-3* exhibited uneven Mst77F staining, suggesting that some nuclei fail to accomplish proper Mst77F incorporation (Supplementary Fig. 6C, D). It is interesting to note that the nuclear decompaction was most prominently observed when males were raised in 25°C after their parents were raised at 18°C (see methods).

Together, these results suggest that *Mst77Y* acts as a dominant negative form of *Mst77F*, interfering with the incorporation of normal protamines and leading to spermatid nuclear decompaction.

## Overexpression of *Mst77Y* results in biased demise in X-bearing spermatids

It has been suggested that rapid copy number expansions on the sex-chromosome likely indicate meiotic drive (*2*). Meiotic drive is expected to cause evolutionary ‘arms race’, wherein competing chromosomes (e.g. X vs. Y) continue to outcompete each other, for example, by increasing the copy number of driver genes. The presence of *Mst77Y* in a multicopy form on Y chromosome implies that *Mst77Y* may favor the transmission of the Y chromosome.

To test the potential involvement of *Mst77Y* in meiotic drive, we examined whether or not X or Y chromosomes are specifically affected by the overexpression of *Mst77Y*. Strikingly, using DNA FISH to distinguish X-vs. Y-bearing spermatids in *Mst77Y*-overexpressing males, we found that decompacting spermatids were more likely to be X-bearing (61.8%) than Y-bearing (38.2%) (Fig. 3F, G). These results suggest that *Mst77Y*, when derepressed, triggers meiotic drive by favoring the viability of Y-bearing sperm. However, the fertility assay revealed only a minor increase in male progeny compared to the background-matched control (51.8% vs. 47.8%, p = 0.0005, Supplementary Fig. 7A), perhaps due to the partial penetrance of spermatid nuclear decompaction upon *Mst77Y* overexpression (Fig. 3E). Nonetheless, clear bias in the nuclear decompaction of spermatids (decompacting nuclei are more likely to contain the X chromosome) indicates that *Mst77Y* acts as a Y-favoring meiotic driver that functions via biased demise of X-chromosome containing spermatids.

Although potentially biased inheritance of sex chromosomes cannot be assessed in the *modulo* mutant (*mod*^*L8*^*/mod*^*07570*^), as the mutant results in eventual demise of all spermatids leading to its sterility, we found that heterozygous *modulo* mutant males (*mod*^*L8*^/+), which are still fertile, exhibited a bias towards male offspring compared to background-matched control (51.4% vs. 46.2%%, p = 0.0001, Supplementary Fig. 7B). This result suggests a possible role for *modulo* in suppressing meiotic drive, perhaps by its ability to suppress the overexpression of *Mst77Y* (Fig. 2C-G).

## Modulo is required to promote polyadenylation of autosomal *Mst77F* transcript

How does *modulo* regulate the expression of *Mst77F* and *Mst77Y*? Modulo protein is expressed in the nucleolus of spermatogonia and spermatocytes, but is downregulated prior to the meiotic divisions (Fig. 4A, B), days earlier than the stages in which its mutant phenotype manifests. Protamine transcripts are also expressed many days prior to the sperm nuclear compaction process (*23*). Interestingly, we found that *Mst77F* transcripts colocalize with Modulo in the spermatocyte nucleolus, while *Mst77Y* transcripts localize adjacent to the nucleolus (Fig. 4C). These results prompted us to examine whether *Mst77F* and/or *Mst77Y* transcripts may be deregulated in *modulo* mutant. Indeed, we found that *Mst77F* mRNA is dramatically reduced in *modulo* mutant, whereas *Mst77Y* mRNA was increased approximately 3-fold using RT-qPCR of polyA-selected RNA (Fig 4D). However, when total RNA was used for RT-qPCR or total RNA sequencing, we found that both *Mst77F* and *Mst77Y* transcripts were increased in *modulo* mutant (Fig. 4D, Supplementary Fig. 8A). RNA FISH, which visualizes total RNA, also confirmed the increase of both *Mst77F* and *Mst77Y* transcripts in *modulo* mutant (Supplementary Fig. 8B). Furthermore, total RNA-seq and RT-qPCR did not detect any splicing defects of *Mst77F* or *Mst77Y* in *modulo* mutant (Supplementary Fig. 10A, B). Collectively, these results suggest that Modulo specifically regulates transcripts of *Mst77F* and *Mst77Y* at the step of polyadenylation.

**Figure 4.**
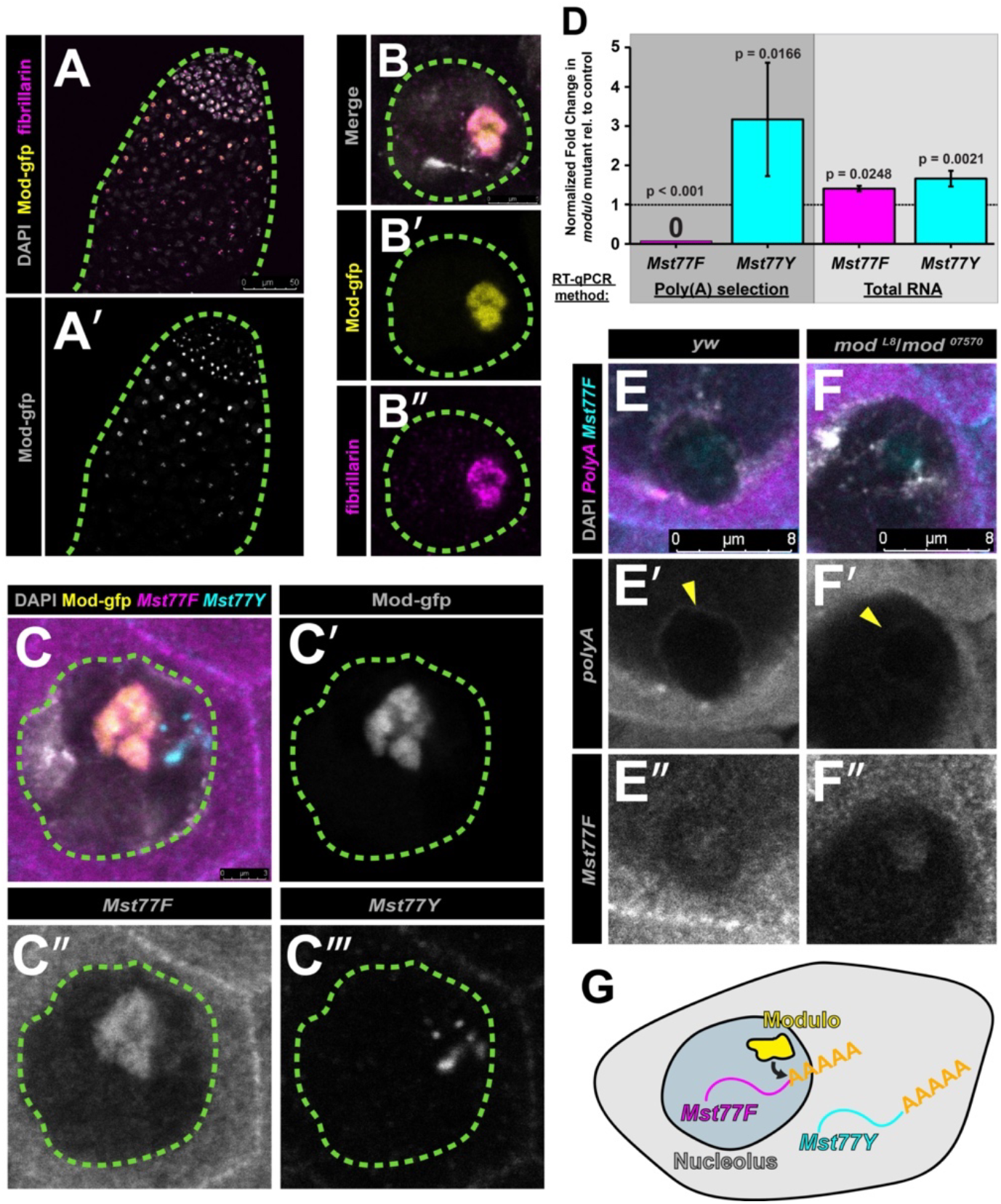
Modulo localizes to nucleolus and functions to promote polyadenylation of *Mst77F*. A, B) Localization of Modulo to the nucleolus in the apical tip of the testis (A) and in the spermatocyte nuclei (B). Modulo-gfp (yellow) stained with anti-fibrillarin (magenta), a nucleolar marker, and DAPI (grey). C) RNA FISH for *Mst77F* and *Mst77Y* transcripts in wild-type spermatocyte nucleus. DAPI (grey), *Mst77F* (magenta), *Mst77Y* (cyan), and Modulo-gfp (yellow). D) RT-qPCR following polyA selection (dark grey) or using total RNA RT-qPCR (light grey) in *modulo* mutant (*mod*^*L8*^/*mod*^*07570*^) vs. sibling control (*mod*^07570^/+) assessing levels of *Mst77F* (magenta) and *Mst77Y* (cyan). Data was normalized to *Rp49* and control. Mean ±SD from three technical replicates is shown. p-values as listed (unpaired Student’s t-test assuming unequal variance on untransformed ddct values). Similar results were obtained from two biological replicates. Primer locations shown in Supplementary Fig. 9a. E, F) RNA FISH for polyA (magenta) and *Mst77F* transcript (cyan) in control(*yw*) (F) vs. *modulo* mutant (*mod*^*L8*^/*mod*^*07570*^) (E), counter-stained with DAPI (grey). *Mst77F* probe was used to identify nucleolus. Yellow arrowhead indicates polyA-containing RNA encircling nucleolus. G) Model for Modulo function in the nucleolus.

Given that Modulo protein and *Mst77F* transcript colocalize in the nucleolus, we speculate that *Mst77F* is directly regulated by Modulo, whereas increased mRNA level of *Mst77Y* may be an indirect consequence of reduced functional *Mst77F* mRNA. Interestingly, RNA FISH using poly(T) probes revealed that poly(A) signal encircles the nucleolus in wildtype spermatocytes, whereas markedly less poly(A) was detected around the nucleolus in the *modulo* mutant (Fig. 4E, F), further suggesting that *modulo* may function to facilitate polyadenylation of transcripts localized to the nucleolus. Taken together, these results suggest that *modulo* plays an essential role in sperm nuclear compaction by facilitating maturation of canonical *Mst77F* transcript over that of Y-linked *Mst77Y* (Fig. 4G).

### Discussion

The present study suggests that *Mst77Y*, a Y-linked multi-copy gene cluster of *Mst77F* homologs, functions as a Y-promoting meiotic driver: we find that *Mst77Y* might act as a dominant-negative protamine variant that preferentially harms the nuclear compaction process of X-bearing spermatids. Interestingly, studies on the Winters sex ratio meiotic drive system in *D. simulans* revealed that the driver *Dox* contains a large portion of the Protamine-like region (*4, 5*). Cytology of the Winters system also suggests failure to properly complete spermiogenesis and condense nuclei (*16*). *Segregation Distorter* (*SD*), a well-studied autosomal drive system in *D. melanogaster* (*1, 3*), was also shown to display spermatid compaction defect with defective incorporation of Protamine A/B (*18*). Importantly, although both *Mst77F/Y* and *Dox* are related to protamines, they are phylogenically distinct from each other^9^. Additionally, *D. simulans* Winters system preferentially harms the Y chromosome, leading to high number of female progeny (*16*), whereas *Mst77Y* preferentially harms the X chromosome-bearing sperm. Moreover, whereas *Dox*-mediated drive is repressed by RNAi, no evidence has been found of small RNA (siRNA or piRNA) for *Mst77Y* (*5, 26*), further suggesting that these drive systems evolved independently. In the absence of RNAi-mediated repression, *modulo*’s role in ensuring polyadenylation of *Mst77F* transcript, while also supporting suppression of *Mst77Y* polyadenylation, may serve as a mechanism to repress meiotic driver(s).

An interesting question that remains is how a dominant negative protamine can preferentially impact a specific chromosome over its homolog. It is possible that such dominant-negative protamines may have preferential affinity to specific chromosomes, perhaps targeting the chromosome-specific DNA sequences and/or chromosome-specific chromatin states. In such a scenario, there must be a sweet spot for the expression level of the ‘dominant-negative, driver protamine’, such that the driver chromosome will incorporate less of the dominant-negative protamine while its opponent will incorporate more. Further upregulation of the dominant-negative protamine would result in the demise of all spermatids, which may contribute to the complete sterility of *modulo* loss-of-function mutant.

Taken together, we propose that a dominant negative form of protamine may be a frequent cause of male meiotic drive by causing a failure to complete nuclear compaction in a subset of gametes containing specific chromosomes. The involvement of protamines in these genetic “arms races” may potentially explain protamines’ known rapid divergence across broad species.

## Methods

### Fly husbandry and strains

All fly stocks were raised on standard Bloomington medium at 25°C, and young flies (1- to 3-d-old adults) were used for all experiments unless otherwise specified. Flies used for wild-type experiments were the standard laboratory wild-type strain *yw* (y^1^w^1^). The following fly stocks were used: *modulo*^*07570*^/TM3 (Bloomington Drosophila Stock Center [BDSC]: 11795), *modulo*^*L8*^/TM3 (BDSC: 38432), and C(1)RM/C(1;Y)6, *y*^*1*^*w*^*1*^*f*^*1*^/0 (BDSC: 9460). The *β2-tubulin* promoter sequence used for producing Mst77Y overexpression was generously provided by Peiwei Chen and Alexei Aravin.

The *Mst77Y* transgenic flies were generated by phiC31 site–directed integration into the Drosophila genome. The *Mst77Y* overexpression sequences in *D. melanogaster* were synthesized by gene synthesis service from Thermo Fisher Scientific (GeneArt Gene Synthesis) and was cloned into pattB vector to insert into specific integration site on 2nd chromosome (attP40) (Supplementary Fig. 5C, Supplementary Table 2). All injection and selection of flies containing integrated transgene were performed by BestGene Inc. (Chino Hills, CA).

Modulo-gfp strain was constructed using CRISPR-mediated knock-in of a gfp-tag at the C-terminus of Modulo (Beijing Fungene Biotechnology Co.) (Supplementary Table 3).

### Sex ratio assay

Individual 1-day old males raised for at least one generation at 18°C were crossed with 3x 1-to 3-day-old virgin *yw* females at 25°C. After 1 day males were removed. This was done to maximize proportion of males exhibiting decompaction phenotype described in Fig. 3. Females were left to produce embryos for a total of 5 days before cleared. Following onset of eclosion, sex of offspring was scored for 10 consecutive days.

### RNA Fluorescent *in situ* hybridization

All solutions used were RNase free. Testes from 1- to 3-day-old flies were dissected in 1X PBS and fixed in 4% formaldehyde in 1X PBS for 30 minutes. Then testes were washed briefly in PBS and permeabilized in 70% ethanol overnight at 4°C. For strains expressing GFP (i.e. Modulo-gfp), the overnight permeabilization in 70% ethanol was omitted. Testes were briefly rinsed with wash buffer (2X saline-sodium citrate (SSC), 10% formamide) and then hybridized overnight at 37°C with fluorescently labeled probes in hybridization buffer (2X SSC, 10% dextran sulfate (sigma, D8906), 1mg/mL E. coli tRNA (sigma, R8759), 2mM Vanadyl Ribonucleoside complex (NEB S142), 0.5% BSA (Ambion, AM2618), 10% formamide). Following hybridization, samples were washed two times in wash buffer for 30 minutes each at 37°C and mounted in VECTASHIELD with DAPI (Vector Labs).

Fluorescently labeled probes were added to the hybridization buffer to a final concentration of 100nM. Poly(T) probes for recognizing Poly(A) sequence were from Integrated DNA Technologies. Probes against *Mst77F* and *Mst77Y* were designed using the Stellaris1 RNA FISH Probe Designer (Biosearch Technologies, Inc.) available online at www.biosearchtech.com/stellarisdesigner. Each set of custom Stellaris1 RNA FISH probes was labeled with Quasar 670 or Quasar 570 (Supplementary Table 4).

Images were acquired using an upright Leica TCS SP8 confocal microscope with a 63X oil immersion objective lens (NA = 1.4) and processed using Adobe Photoshop and ImageJ software.

### DNA Florescent *in situ* hybridization

Testes from 1–3-day old flies were rapidly dissected in 4% formaldehyde, 1mM EDTA in 1X PBS then nutated for 30 minutes. Then testes were washed three times in 1X PBST (PBS containing 0.1% Triton-X) +1mM EDTA for 30 minutes each. Testes were briefly rinsed with 1X PBST and then incubated at 37°C for 10 minutes with 2 mg/mL RNase A in PBST. Following Rnase treatment, samples were washed once in 1X PBST + 1mM EDTA for 10 minutes. Samples were then briefly rinsed with 2X SSC + 1mM EDTA + 0.1% Tween-20, and then washed three times in 2X SSC + 0.1% Tween-20 + formamide (20% for first wash, 40% for second, 50% for third) for 15 minutes each. Samples were washed then with 2X SSC + 0.1% Tween-20 + 50% formamide. Samples were then incubated for 5 minutes at 95°C with fluorescently labeled probes in hybridization buffer (2X SSC, 10% dextran sulfate, 50% formamide, 1mM EDTA) then transferred to 37°C overnight. Following hybridization, samples were washed three times in 2X SSC + 1mM EDTA + 0.1% Tween-20 for 20 minutes each then mounted in VECTASHIELD with DAPI (Vector Labs).

Fluorescently labeled probes were added to the hybridization buffer to a final concentration of 500nM. Satellite DNA probes distinguishing X and Y chromosomes (AATAC)_6_-Cy5 for Y and (TAGA)_8_-Cy3 were from Integrated DNA Technologies.

### Immunofluorescence staining (IF)

Testes were dissected in 1X PBS, transferred to 4% formaldehyde in 1X PBS, and fixed for 30 min. Testes were then washed three times in 1X PBST (PBS containing 0.1% Triton X-100) for 20 minutes each followed by incubation with primary antibodies diluted in 1X PBST with 3% BSA at 4°C overnight. Samples were washed three times in 1X PBST for 20 minutes each, incubated with secondary antibody in 1X PBST with 3% BSA for 2 hours at room temperature, Samples were then washed three times in 1X PBST for 20 minutes each and mounted in VECTASHIELD with DAPI (Vector Labs).

The following primary antibodies were used: anti-fibrillarin (1:200, mouse; Abcam; ab5812), anti-Modulo (1:1000, guinea pig; this study), anti-Protamine A/B (1:200, guinea pig, gift of Elaine Dunleavy, Centre for Chromosome Biology, National University of Ireland, Galway, Ireland (*27*), anti-Mst77F (1:1000; guinea pig, this study), anti-Mst77Y (1:500; rabbit, this study), anti-Tpl94d (1:500; rabbit, this study), Phalloidin-Alexa Fluor 546 or 488 (1:200; Thermo Fisher Scientific; A22283 or A12379). The Modulo antibody was generated by injecting a peptide sequence CRKQPVKEVPQFSEED[48-62] targeting the N-terminal end of Modulo) in guinea pigs (Covance). The Tpl94d antibody was generated by injecting a peptide DKGSAYKPLTLNRSYVIRKC[96-114] in rabbits (Covance). The Mst77F antibody was generated by injecting multiple peptides (SKPEVAVTC[9-16], YKKSIEYVNC[22-30], CRSSEGEHR[112-119], LQRSSEGEHRMHSEC[110-123], RSSGKPKPKGARPRKC[169-183]) targeting sites in Mst77F, as indicated, differentiating it from Mst77Y in guinea pigs (Covance). The Mst77Y antibody was generated by injecting multiple peptides (IKPDVAVSC[9-16], SRKAIEYVKC[22-30], CRSIEAELR[112-119], KTSRKAIEYVKSD[20-32], CVSSLQRSIEAELR[107-119]) targeting sites of varying aa length in Mst77Y differentiating it from Mst77F in rabbits (Covance). Alexa Fluor–conjugated secondary antibodies (Life Technologies) were used at a dilution of 1:200.

### RT-qPCR

Total RNA was purified from *Drosophila melanogaster* adult testes (75 pairs/sample) by Direct-zol RNA Miniprep (Zymo Research), with DNase treatment according to manufacturer’s protocol. 1μg of total RNA was reverse transcribed following priming with either random hexamers or polyT using SuperScript III® Reverse Transcriptase (Invitrogen) followed by qPCR using Power SYBR Green reagent (Applied Biosystems). Primers for qPCR were designed to amplify mRNA or intron-containing transcript as indicated. Relative expression levels were normalized to Rp49 and control siblings. All reactions were done in technical triplicates with at least two biological replicates. Graphical representation was inclusive of all replicates and p-values were calculated using a t-test performed on untransformed average ddct values. Primers used are described in Supplementary Fig. 9A, B.

### Total RNA-seq

Total RNA was purified from *Drosophila melanogaster* adult testes by Direct-zol RNA Miniprep (Zymo Research), with Dnase treatment. Quality of indexed libraries was confirmed using an Agilent Fragment Analyzer and qPCR. Sequencing libraries were prepared with the KAPA RNA HyperPrep Kit with RiboErase. Samples were sequenced on a HiSeq 2500, producing 100×100 nt paired-end reads. The read pairs were mapped to the canonical chromosomes of the Drosophila melanogaster genome (assembly BDGP6/dm6) using STAR 2.7.1a (*28*); default parameters, except ‘—alignIntronMax 25000’, indexed with all FlyBase genes (FB2020_06 Dmel Release 6.37) and the option ‘—sjdbOverhang 100’. Gene counts were obtained using featureCounts (*29*); v 2.0.1, with ‘-M –fraction -p -s 2’). After summing gene counts for technical replicates, differential expression was assayed using DESeq2 (*30*) v1.26.0, with lfcShrink(type=“ashr”)). RNA coverage across genes at nucleotide resolution was quantified with “bedtools coverage” (*31*) and scaled by the total number of reads mapped to genes.

### Statistics and reproducibility

Data are presented as mean ± s.d. unless otherwise indicated. The sample number (n) indicates the number of testes, nuclei, or male flies in each experiment as specified in the figure legends. We utilized two-sided Student’s t-tests to compare paired or independent samples, as applicable and is specified in the figure legends. We calculated probability using exact binomial distribution with parameters specified in Fig. 3G legend. No statistical methods were used to predetermine sample sizes. The experimenters were not blinded to the experimental conditions, and no randomization was performed. All the statistical details of the experiments are provided in the main text and legends. P-values listed either in figure, figure legends, or Supplementary Table 1.

### Reporting summary

Further information on research design is available in the Nature Research Reporting Summary linked to this article.

### Data availability

Sequencing data is available at NCBI GEO under accession GSE214456. All other data needed to evaluate the conclusions in the paper are present in the paper and/or the Supplementary materials.

## Acknowledgments

We thank the Bloomington Drosophila Stock Center, Dr. Elaine Dunleavy for reagents. We thank the Data Science, Bioinformatics, and Informatics Core at the University of Michigan for consulting, Dr. Bing Ye for advice and support. We thank the Yamashita, Lehmann, and Ye lab members, Drs. Daven Presgraves and Eric Lai for discussions and Yamashita Lab members for comments on the manuscript. The research was supported by the NIH (to J.I.P., F30HD105324), Howard Hughes Medical Institute (to Y.M.Y.), and Whitehead Institute for Biological Research.

## Author information

### Authors and Affiliations

**Life Sciences Institute, University of Michigan, Ann Arbor, Michigan, USA**

Jun I. Park

**Whitehead Institute, Massachusetts Institute of Technology, Cambridge, Massachusetts, USA**

George W. Bell, Yukiko M. Yamashita

## Contributions

Conceptualization: JIP, YMY

Methodology: JIP, GWB, YMY

Investigation: JIP, GWB

Visualization: JIP, GWB, YMY

Funding acquisition: JIP, YMY

Project administration: JIP, YMY

Supervision: YMY

Writing – JIP, YMY

Writing – review & editing: JIP, GWB, YMY

## Ethics Declarations

Competing interests

The authors declare no competing interests

## Supplementary figures and tables

**Supplementary Figure 1.**
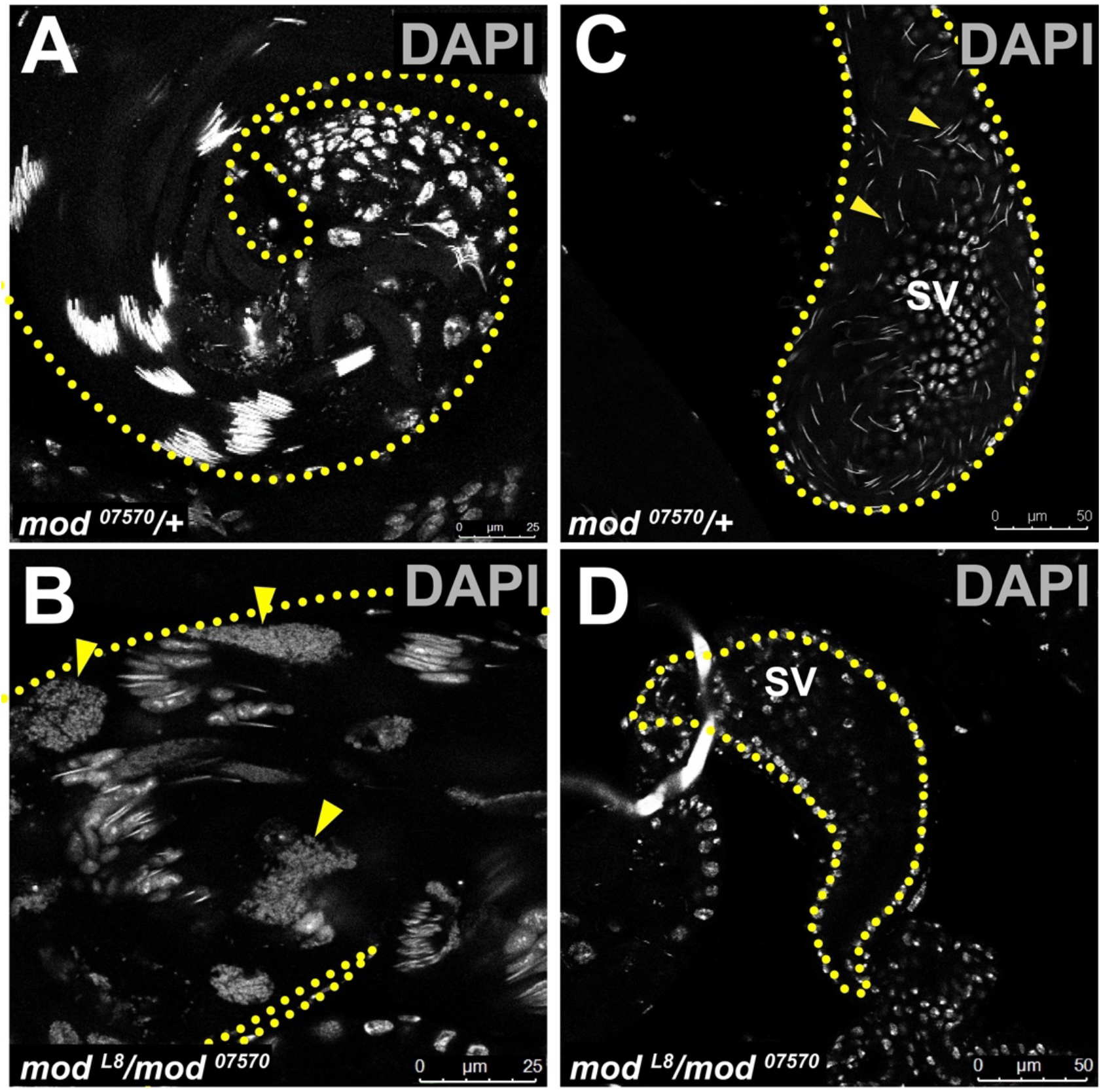
*Modulo* mutant results in sterility with complete spermatid demise and empty seminal vesicles. A, B) Representative images of the testis basal end of control (*mod*^*07570*^/+) (A) and mutant (*mod*^*L8*^/*mod*^*07570*^) (B) males. Arrowheads pointing to widespread DAPI-positive debris present in *modulo* mutant. DAPI (grey). Dotted line outlines the testis basal end (A, B) C, D) Representative images of seminal vesicle of control (*mod*^*07570*^/+) (C) and mutant (*mod*^*L8*^/*mod*^*07570*^) (D) males. ‘SV’ indicating seminal vesicle. DAPI (grey). Dotted line outlines the seminal vesicle (C, D).

**Supplementary Figure 2.**
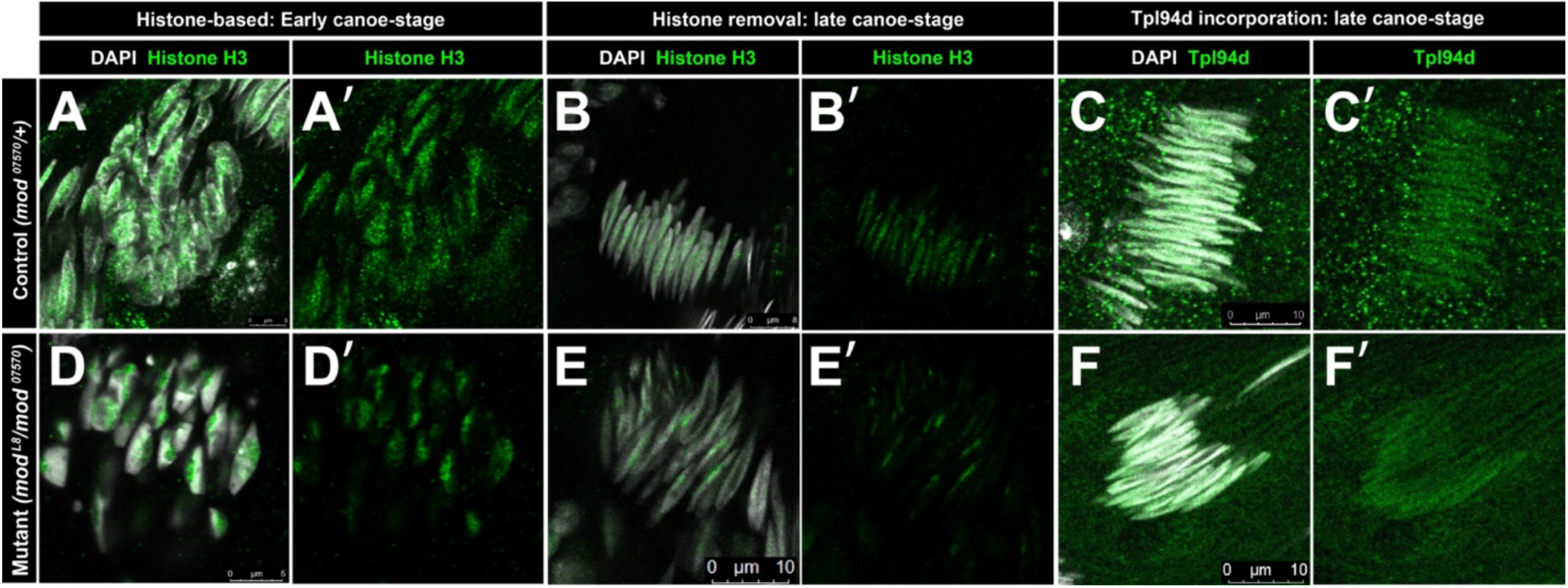
*Modulo* mutant proceeds normally through early stages of histone-to-protamine transition. A-F) Representative images of control (*mod*^07570^/+) (A-C) and mutant (*mod*^*L8*^/*mod*^07570^) (D-F) spermatids undergoing appropriate histone removal and transition protein incorporation.

**Supplementary Figure 3.**
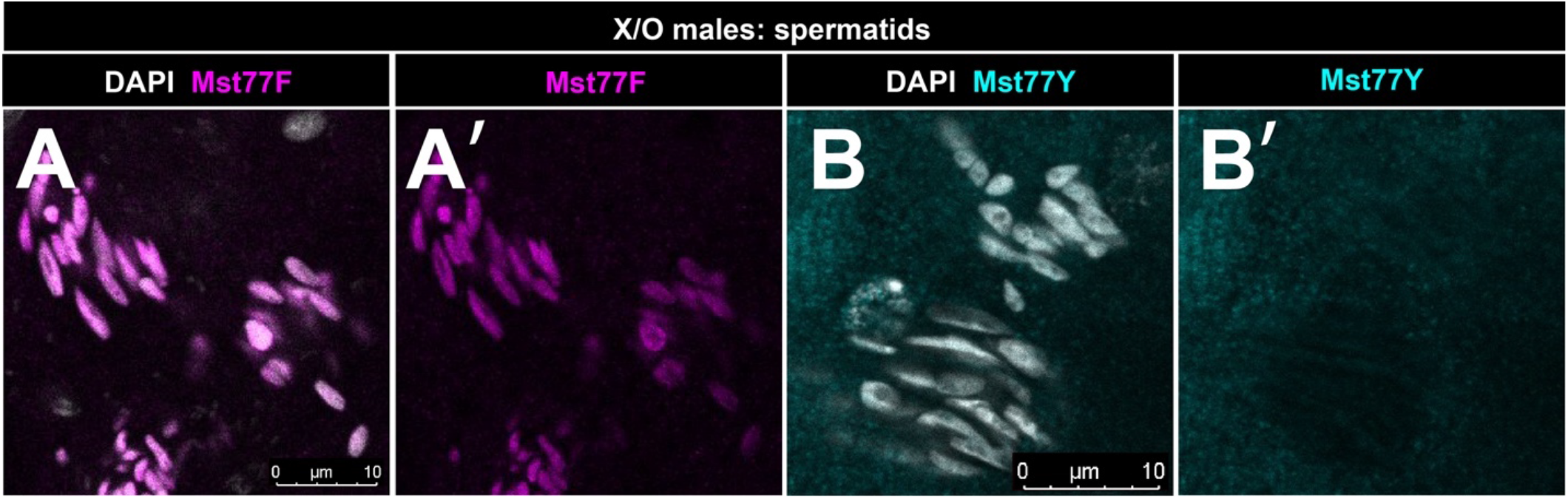
Specificity of Mst77Y antibody. A) Representative image of IF using anti-Mst77F in XO males, showing expected presence of Mst77F, an autosomal gene. B) Representative image of IF using anti-Mst77Y in XO males showing expected absence of Mst77Y, a Y-chromosome gene.

**Supplementary Figure 4.**
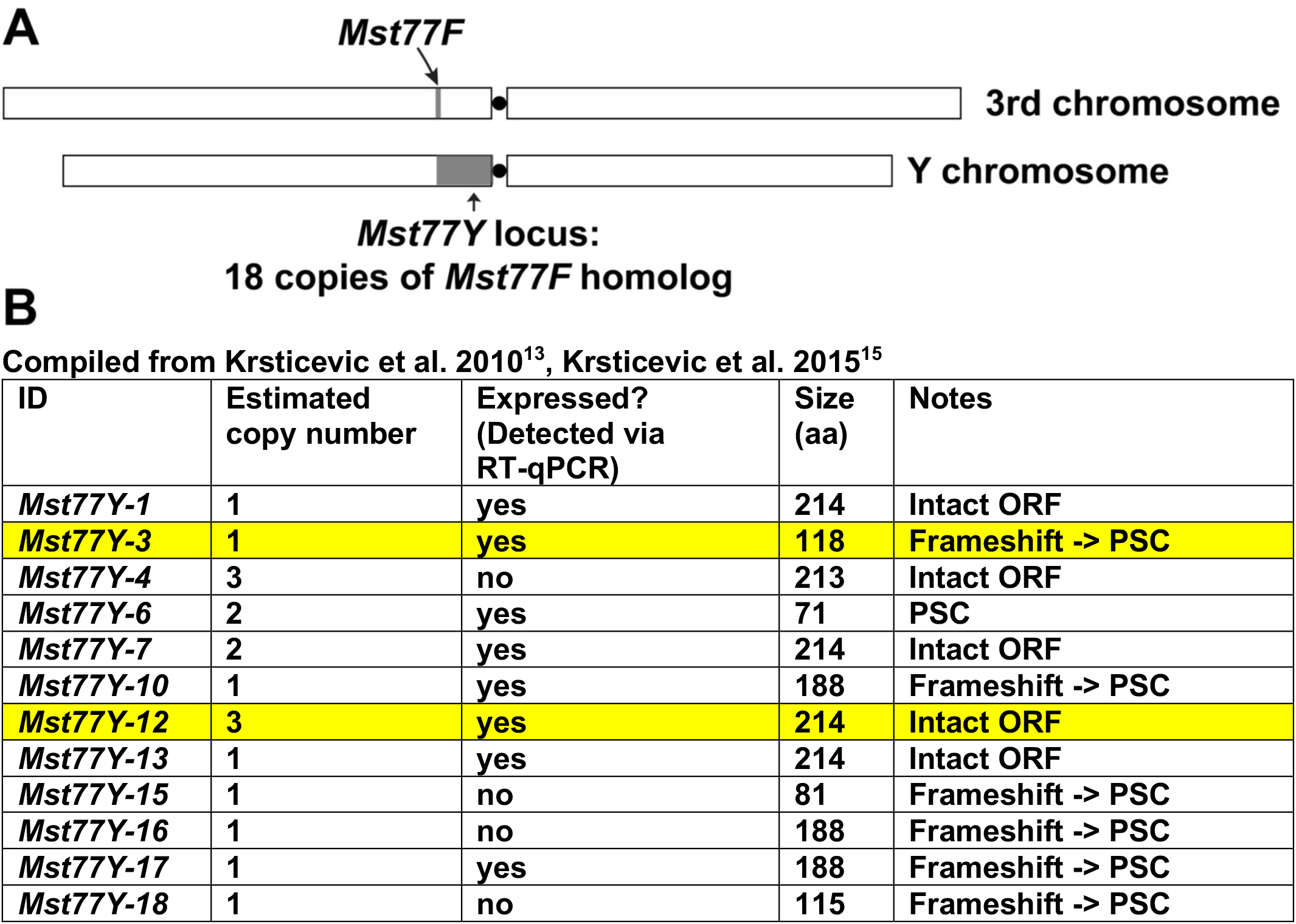
*Mst77Y* locus contains 18 copies of *Mst77F* homolog located on the Y-chromosome. A) Location of *Mst77Y* and *Mst77F* loci. Shaded regions are not drawn to scale. B) Summary of *Mst77Y* genes (18 copies of *Mst77F* homologs), compiled from Krsticevic et al. 2010^16^ and Krsticevic et al. 2015^17^. Yellow highlighted rows indicate the genes used in this paper for overexpression in transgenic animals.

**Supplementary Figure 5.**
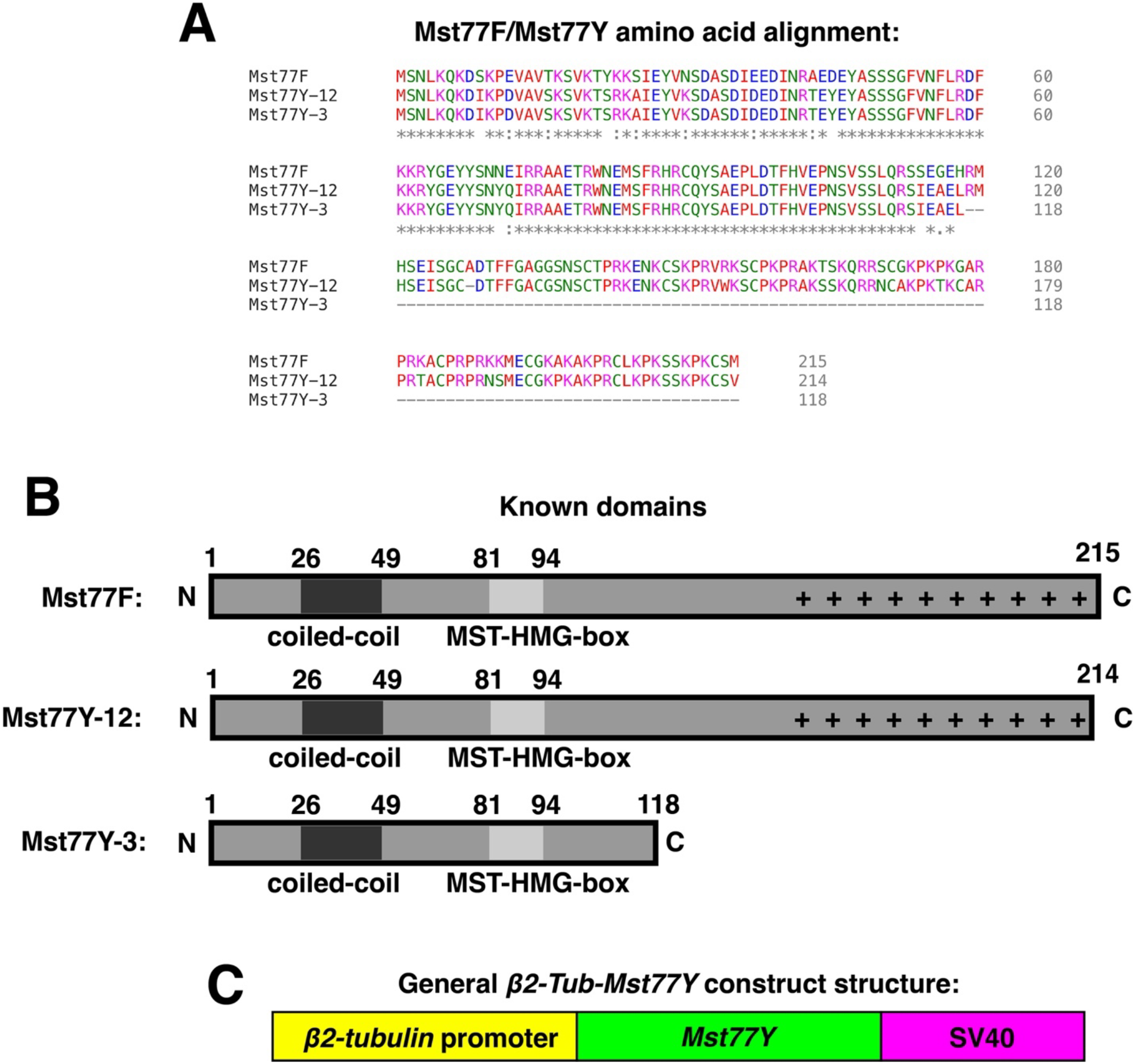
Schematic of *β2-tub*-Mst77Y constructs. A) Amino acid sequence alignment of Mst77F, Mst77Y-3, Mst77Y-12. Asterisks (*) represent identical residue, colons (:) represent non-identical but similar amino acids, blank space indicates non-identical and non-similar amino acids, and hyphens (-) indicate missing residues. B) Schematic of previously described Mst77F domains based on Doyen et al., 2015 (*32*) and Kost et al., 2015 (*33*). “+” indicates region of highly positively charged residues. C) Structure of transgenic construct, consisting of *β2-tubulin* promoter followed by *Mst77Y* ORF (either *Mst77Y-12 or Mst77Y-3*), stop codon and SV40 3’-UTR. Sequence information contained in Supplementary Table 2.

**Supplementary Figure 6.**
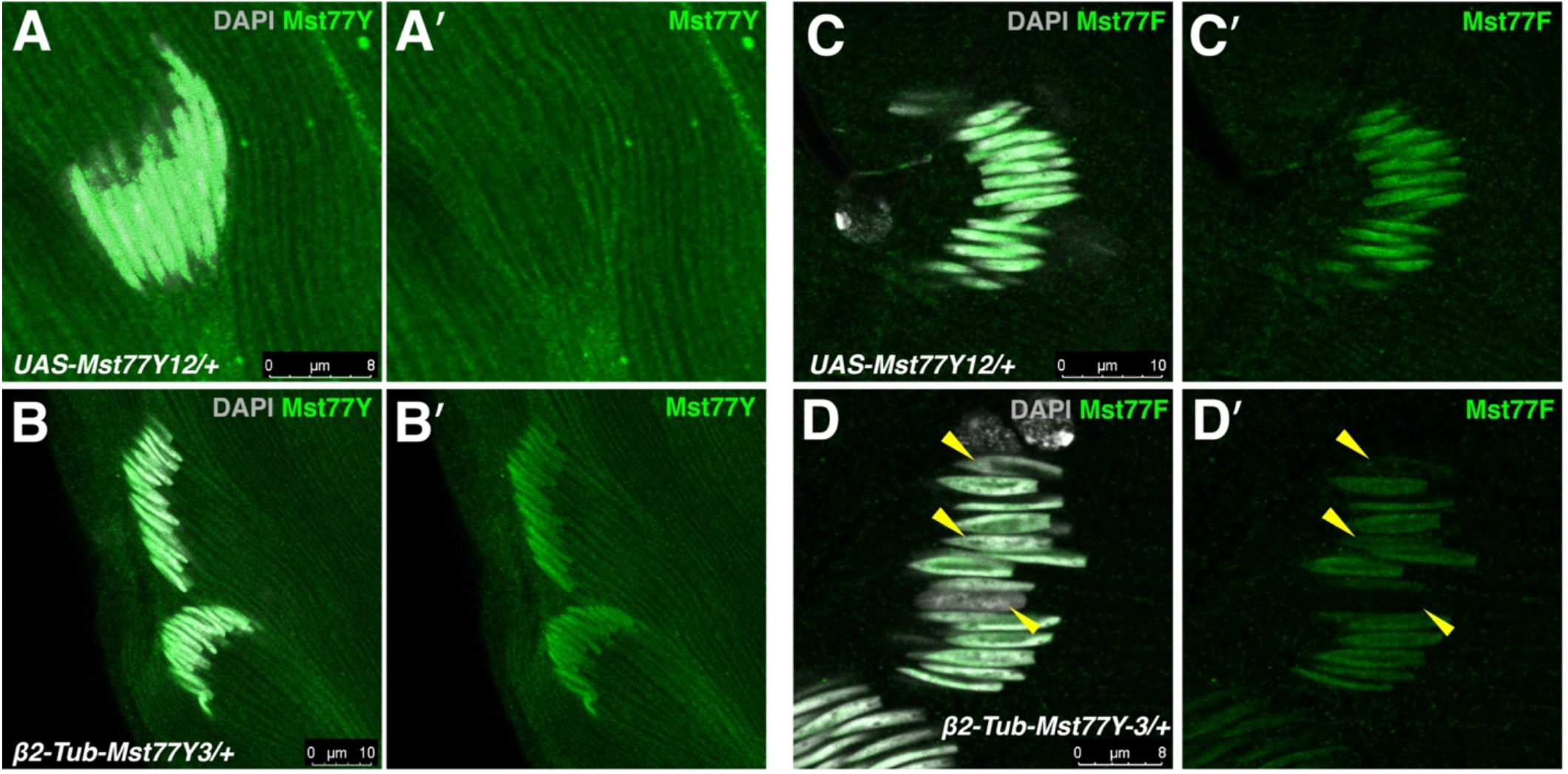
Transgenic *Mst77Y* flies overexpress Mst77Y and display Mst77F incorporation defects. A, B) Representative images of IF using anti-Mst77Y in control (*UAS-Mst77Y12*/+, without gal4 driver) (A) and *β2-tub-Mst77Y3*/+ (B). C, D) Representative images of IF using anti-Mst77F in *UAS-Mst77Y12/+* (C) and *β2-tub-Mst77Y3* (D) males. Yellow arrowheads indicate spermatid nuclei within the same cyst with decreased Mst77F incorporation.

**Supplementary Figure 7.**
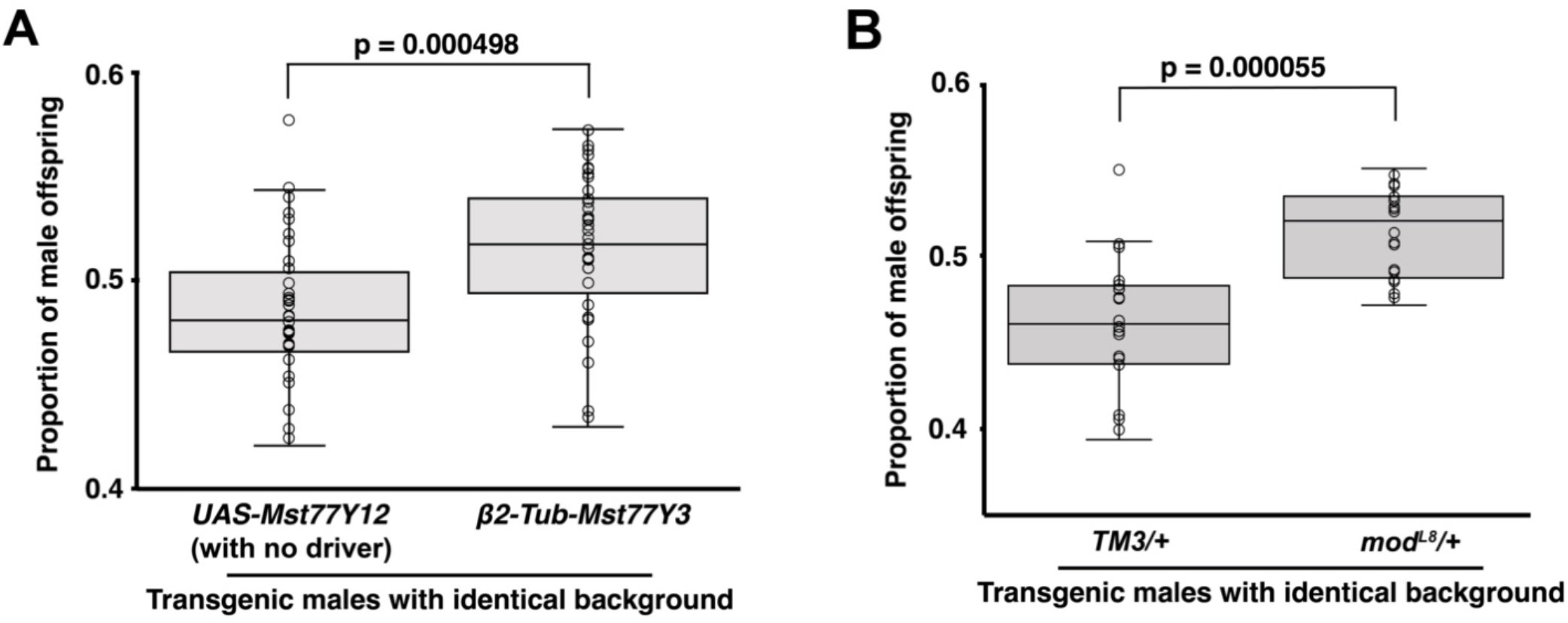
*Modulo* mutant heterozygote males and Mst77Y overexpressing males display male-biased sex ratio. Box and whisker plot showing proportion of male offspring produced by transgenic males in a sex ratio assay. A) Control males (*UAS-Mst77Y12*/+) with no driver or Mst77Y overexpressing males (*β2-tub*-*Mst77Y*3/+) were crossed with 3 *yw* virgin females and sex of resultant offspring were scored. p-value shown in figure (unpaired Student’s t-test assuming unequal variance) with n = 35 individual crosses in control and n = 36 individual crosses in experimental condition from 3 independent experiments. B) Control males (*TM3/+*) or *modulo* mutant heterozygote males (*mod*^*L8*^/+) were crossed with 3 *yw* virgin females and sex of resultant offspring were scored. p-value shown in figure (unpaired Student’s t-test assuming unequal variance) with n = 19 individual crosses in control and n = 18 individual crosses in experimental condition from 3 independent experiments.

**Supplementary Figure 8.**
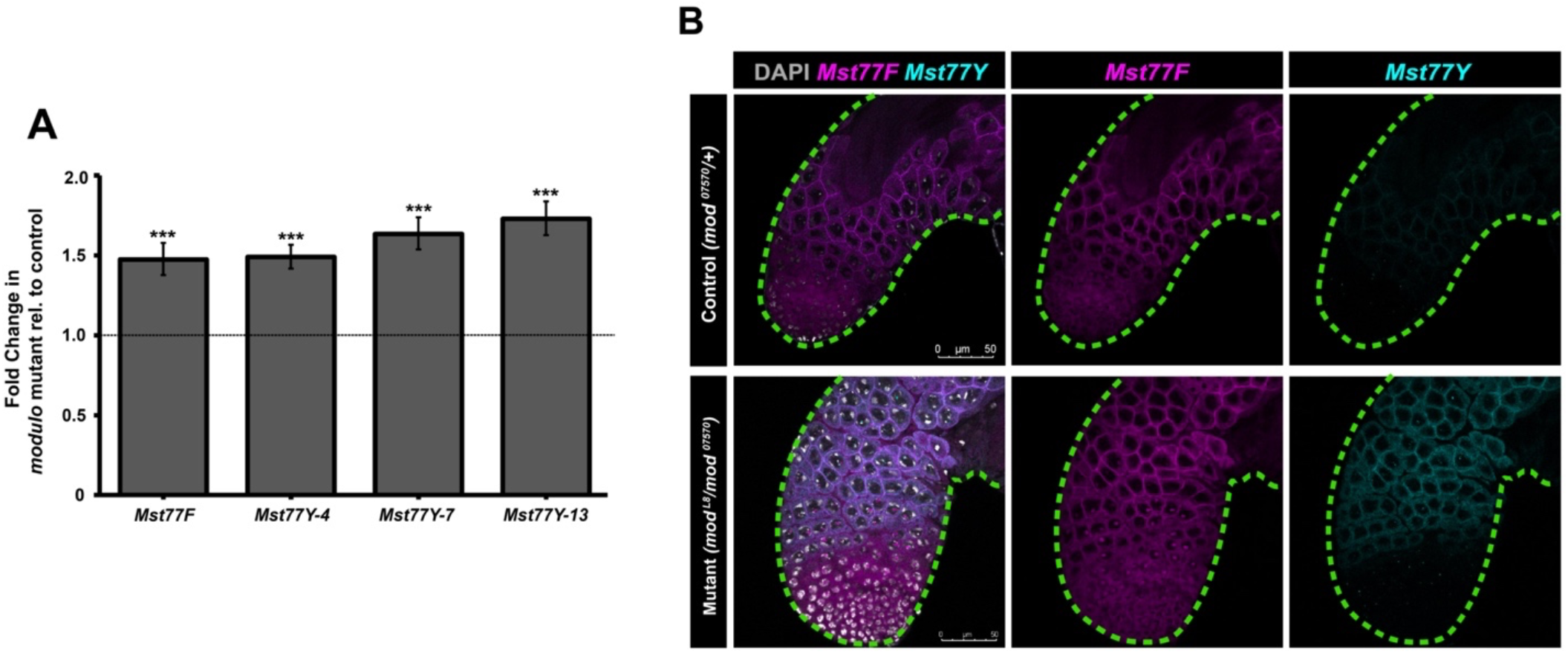
*Modulo* mutant results in up-regulation of *Mst77F* and *Mst77Y* total RNA. A) Fold-change of *Mst77F* and *Mst77Y* RNA in *modulo* mutant (*mod*^L8^/*mod*^*07570*^) compared to control (*mod*^07570^/+) using total RNA-seq. *** indicates p < 0.001 (unpaired Student’s t-test assuming unequal variance) with n = 3 independent experiments. B) Representative images of RNA FISH of *Mst77F* (magenta) and *Mst77Y* (cyan) in control (*mod*^07570^/+) and *modulo* mutant (*mod*^L8^/*mod*^07570^) males.

**Supplementary Figure 9.**
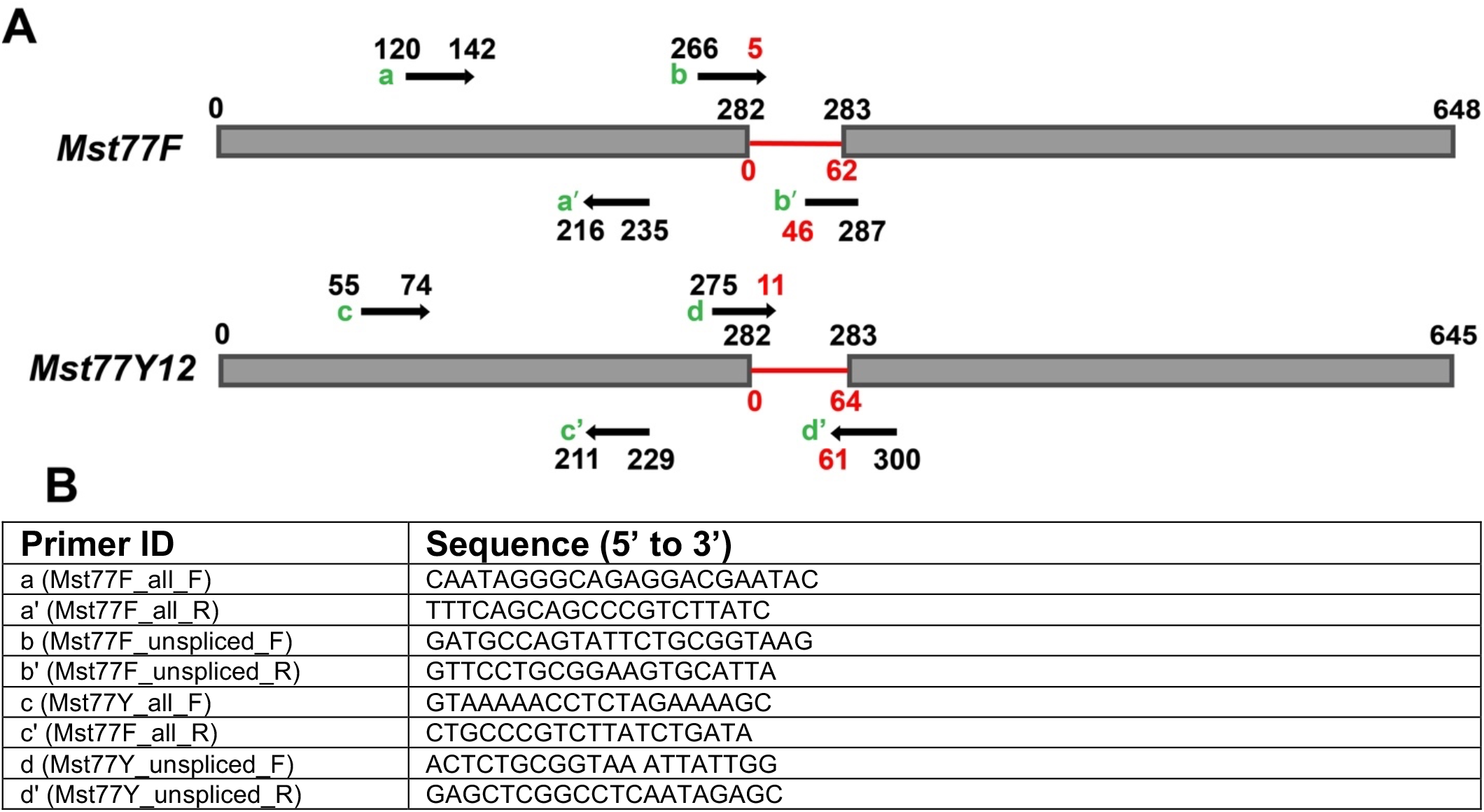
RT-qPCR primers for *Mst77F/Mst77Y*. A) schematic of *Mst77F* and *Mst77Y* primer sets for RT-qPCR relative to location of exon junctions of respective genes. B) table of respective primer sequences. Primers specific to *Mst77F* or *Mst77Y* could not be designed to span exon-exon junction due to high sequence homology in this region. Splicing changes were assessed by looking at relative amount of total transcript level vs. unspliced transcript level.

**Supplementary Figure 10.**
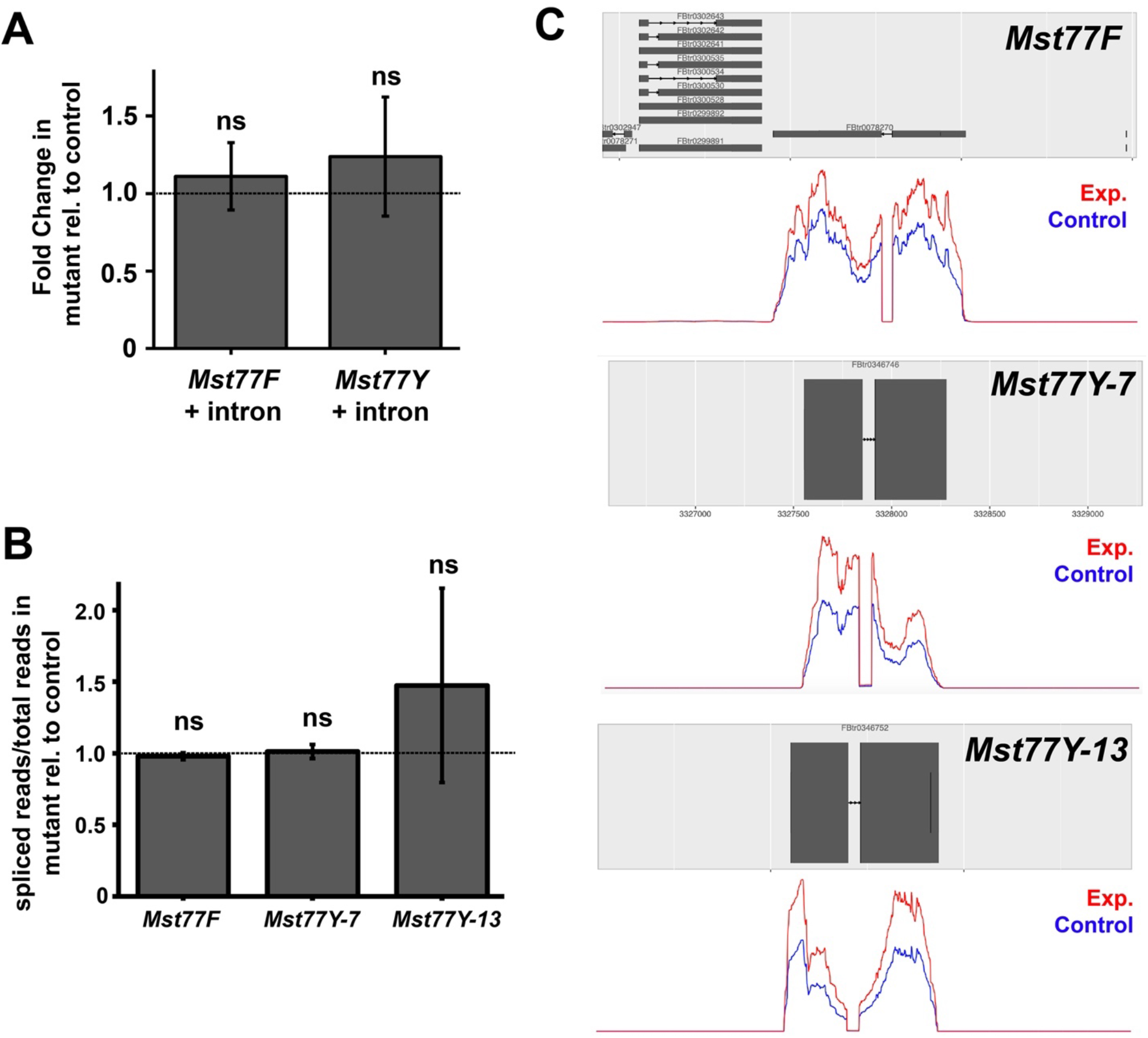
*Modulo* mutant does not affect splicing of *Mst77F* and *Mst77Y* transcript. A) RT-qPCR using exon-intron primer sets to examine level of nascent intron-containing transcript of *Mst77F* and *Mst77Y* in mutant (*mod*^*L8*^/*mod*^*07570*^) compared to control (*mod*^*07570*^/+). Primer sets provided in Supplementary Fig. 9. Data was normalized to Rp49 and control. Mean ±SD from three technical replicates is shown. p-values calculated using unpaired Student’s t-test assuming unequal variances with n = 6 replicates from 2 independent experiments. Similar results were obtained from two biological replicates. Primer locations shown in Supplementary Fig. 9A. B) splicing analysis using total RNA-seq reads comparing proportion of spliced reads to total reads in mutant (*mod*^*L8*^/*mod*^*07570*^) normalized to control (*mod*^*07570*^/+). p-values calculated using unpaired Student’s t-test assuming unequal variances with n = 3 independent experiments. C) Scaled read coverage of *Mst77F, Mst77Y-7, Mst77Y-13* genes showing similar coverage between mutant (*mod*^*L8*^/*mod*^*07570*^) and control (*mod*^*07570*^/+) of exons/introns *Mst77F* and *Mst77Y* genes.

## Supplementary Information

**Supplementary Table 1.** P-values in this study.

| Figure label | p-value |
| --- | --- |
| Fig. 1K | 0 |
| Fig. 2G | $7.188 \times 10^{-7}$ |
| Fig. 3E | $7.34 \times 10^{-4}$ (*UAS-Mst77Y12/+* vs. *β2-tub-Mst77Y3/+*)<br>$3.64 \times 10^{-3}$ (*UAS-Mst77Y12/+* vs. *β2-tub-Mst77Y12/+*)<br>0.750 (*β2-tub-Mst77Y3/+* vs. *β2-tub-Mst77Y12/+*) |
| Fig. 3G | $1.18 \times 10^{-4}$ |

**Supplementary Table 2.**
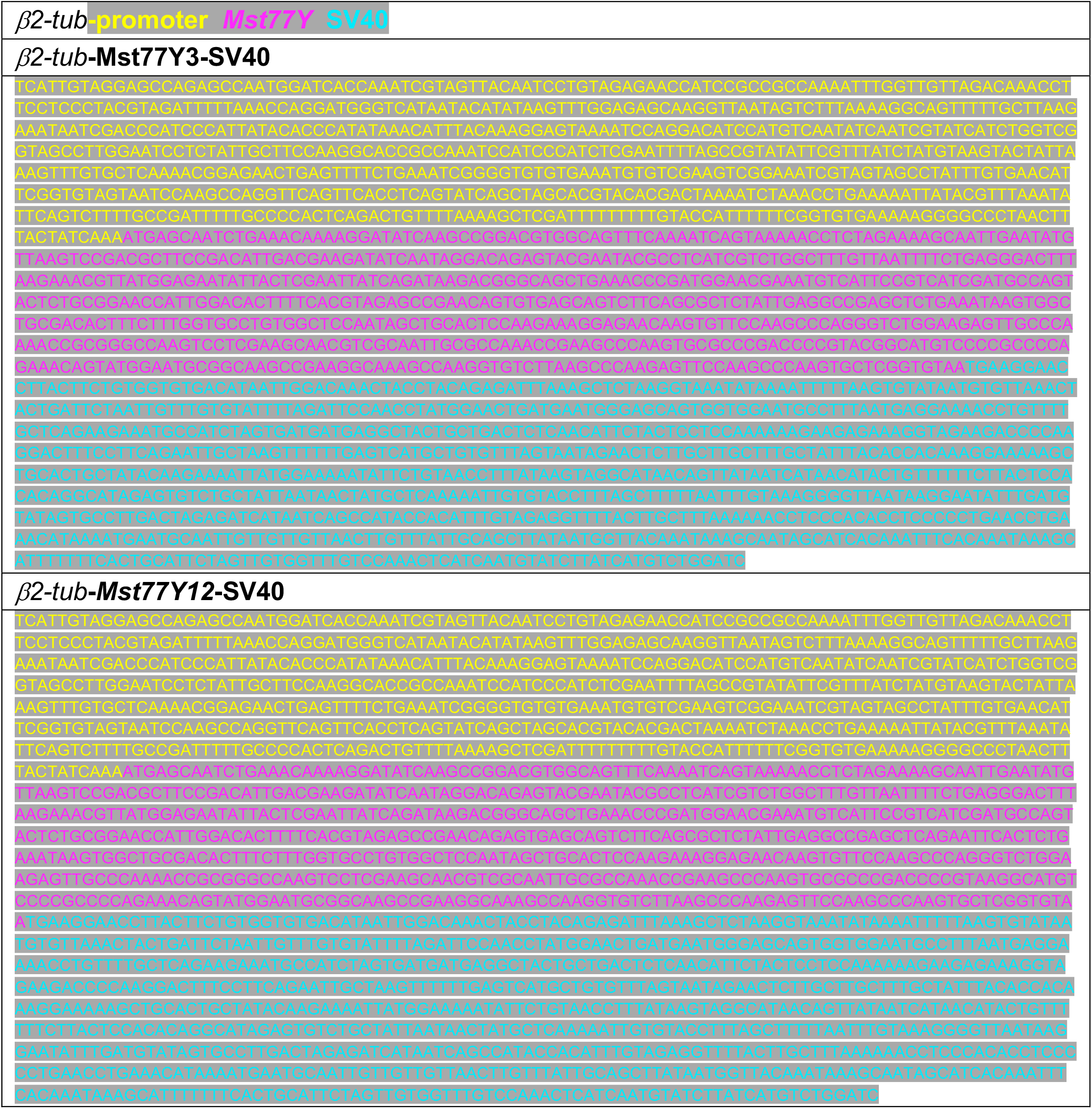
Mst77Y constructs utilized to generate transgenic Mst77Y overexpression lines. *β2-tub* promoter sequence (yellow), respective Mst77Y sequence (magenta), SV40 3’ UTR sequence (cyan)

**Supplementary Table 3.**
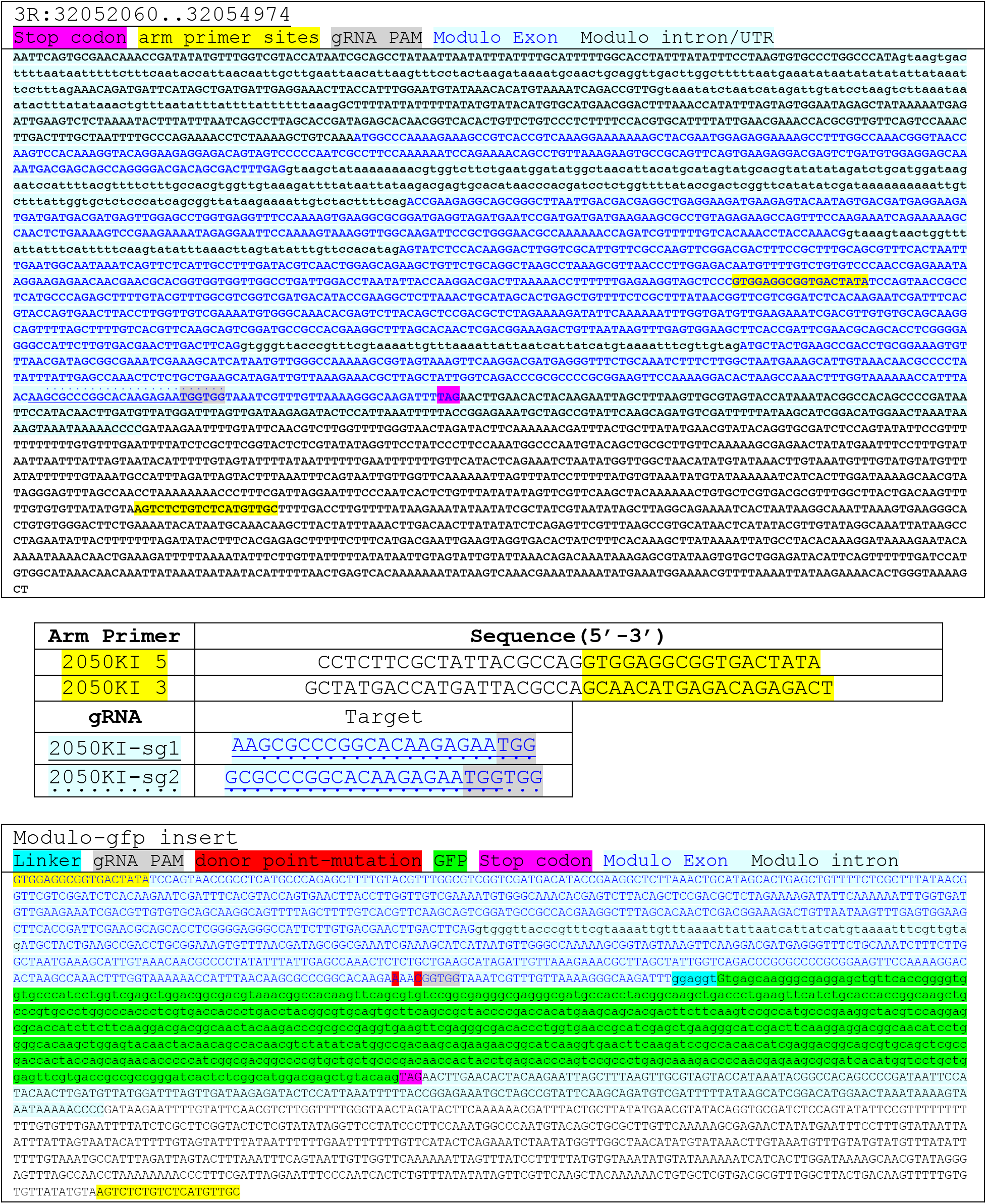
CRISPR/Cas9 generation of *modulo-gfp* at endogenous locus.

**Supplementary Table 4.**
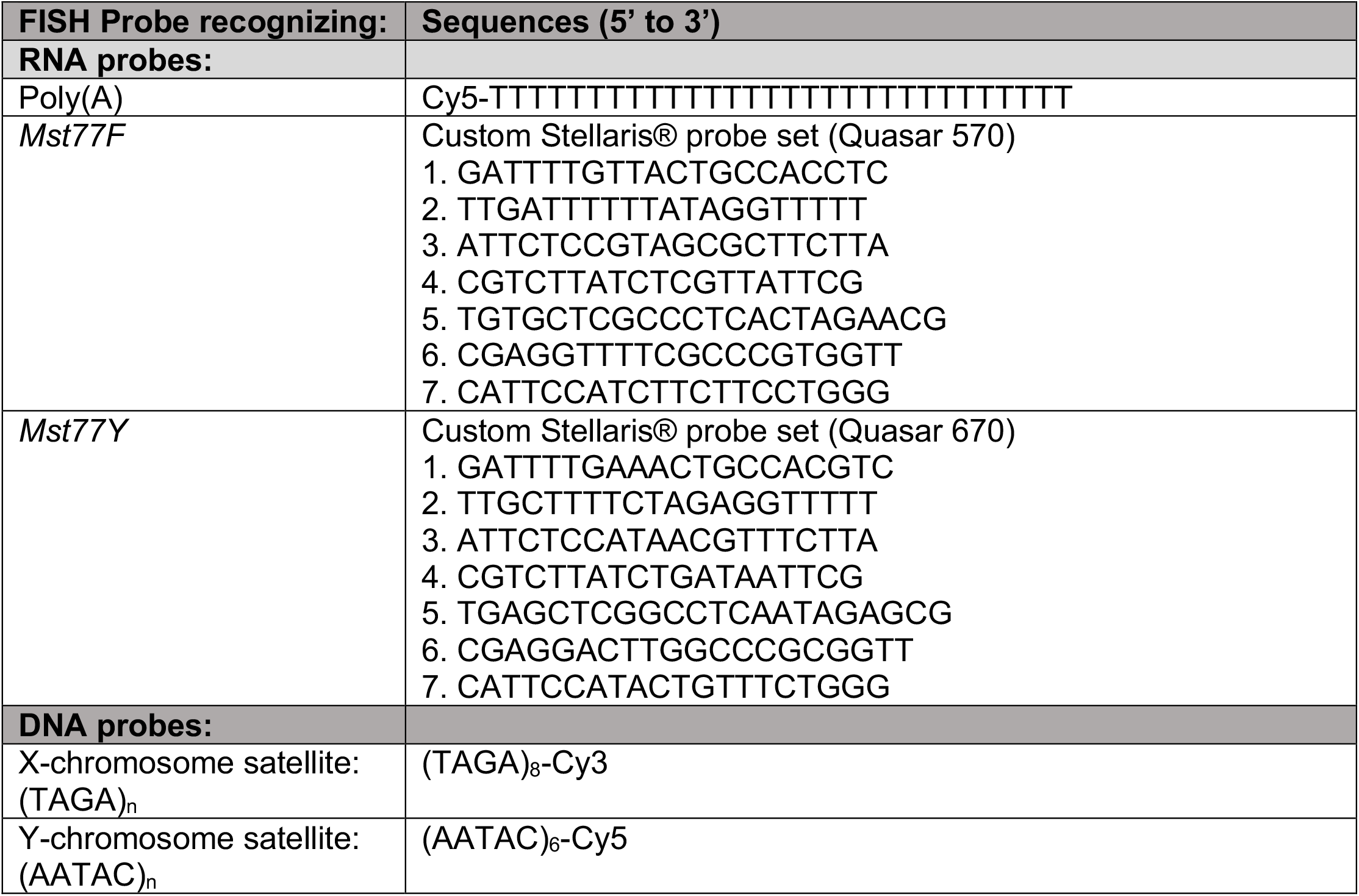
FISH probes utilized in this study.

